# Direct fluorescence detection and volume electron microscopy reveal a role for antibiotic biosynthesis in the bacterial cell envelope of *Streptomyces sp.* Mg1

**DOI:** 10.64898/2026.01.23.701342

**Authors:** Nitya Kalyani Josyula, Anton Classen, Alma Fernandez, Aart J. Verhoef, Anindito Sen, Paul D Straight

## Abstract

Secondary metabolites support the environmental fitness of a broad taxonomic diversity of bacteria and fungi, yet the cellular biology that supports secondary metabolism is little understood. This study focuses on the linearmycin antibiotics from *Streptomyces sp*. Mg1. These membrane-disruptive metabolites are packaged into extracellular vesicles, which in turn require the linearmycins for their biogenesis. This connection suggests that for some secondary metabolites, their biosynthesis is an integral function of cell organization and physiology. In this study, we employed simultaneous multi-photon fluorescence microscopy to directly detect linearmycins, enabling us to follow their biosynthesis and accumulation. We found that linearmycin fluorescence localizes to membrane-dense regions that also coincide with localization with a fluorescent fusion protein, LnyI-Ypet, that is essential for biosynthesis. Genetic disruption of linearmycin biosynthesis led to changes in the cell membrane visible using lipophilic dyes. To resolve differences between wild type and linearmycin-deficient membranes, we used serial FIB milling and scanning electron microscopy (FIB-SEM) to generate 3D volume reconstructions of *S*. Mg1 filaments. Using this approach, we identified granular subcellular compartments that require linearmycins. Disruption of linearmycin synthesis causes, in addition to disappearance of the compartments, visible distortions in the cell envelope, suggesting an integral role for the metabolites in membrane dynamics of the producer bacteria. We propose that the subcellular compartments coalesce near hyphal cell division and branch points and are regions for linearmycin biosynthesis. This study demonstrates the combined use of advanced microscopy to reveal an intrinsic role for antibiotics in the cell biology of the producing organism.

## Introduction

Secondary metabolites are key mediators of microbial competition in the environment (1,2). Actinomycetes are responsible for the production of > 60% of bioactive metabolites that are considered medically and pharmaceutically relevant (3–5). While much research focuses on therapeutics and their mechanisms of action, the pertinence of these metabolites for the producing organisms is less explored. The production of many secondary metabolites is life-cycle dependent, suggesting their biosynthesis is entrained to the physiology of the producing bacteria. For example, prodigiosins are pigmented tripyrrole compounds produced by *Streptomyces coelicolor* (6,7). They accumulate in mycelia late in the life cycle and trigger programmed cell death that results in release of the metabolite (8). The linearmycins are a family of polyene antibiotics that are packaged into extracellular vesicles by *Streptomyces sp*. Mg1 and cause lysis of *B. subtilis* by damaging the membrane (9). The biosynthesis of the linearmycins is required for extracellular vesicle biogenesis, suggesting that antibiotic biosynthesis and membrane biogenesis may be integrated processes. In order to determine how these processes connect, we sought imaging approaches to support in-depth biochemical and genetic studies.

Filamentous bacteria present multiple technical challenges for imaging cellular processes. These include the small size of EVs relative to cells, and the 3D networks of branched hyphae that constitute mycelia. The use of classical fluorescent reporters for protein localization in *Streptomyces* is difficult due to high background fluorescence within the hyphae (10). Label-free imaging approaches provide new tools for microscopy of filamentous organisms. Imaging mass spectrometry has proven useful to determine spatial distribution of metabolites (11–14). Similarly, Raman spectroscopy was used to detect amphotericin B and its association with the mycelia of *Streptomyces nodosus* (15). Autofluorescence of polyene antibiotics is an effective tool to visualize membrane cell biology in eukaryotic cells (16–19). We demonstrated the use of simultaneous two- and three-photon fluorescence imaging to detect autofluorescence of a polyene macrolide, ECO-02301, from *Streptomyces aizunensis,* resolving single filaments up to 750 μm depth within a mycelium (20). Higher-resolution techniques, especially electron microscopy (EM), provide additional approaches to visualize cells and subcellular organization. SEM of *Streptomyces* has been used to determine surface and spore morphology (21,22) while cryo-electron tomography (ET) has enabled visualization of vegetative growth, sporulation and spore structure without damaging the cells (23–26). Recently, (cryo)-FIB-SEM has emerged as a valuable tool for volume EM to visualize nanoscale cellular features in three dimensions (27).

In this study, we employed a combination of two and three photon fluorescence and electron microscopy to detect and localize linearmycins and their potential sites of biosynthesis in whole *S.* sp. Mg1 cells. These approaches enabled us to image the cellular distribution of linearmycins and to identify phenotypes associated with their loss. Direct detection by three-photon excitation fluorescence microscopy revealed subpopulations of *S*. Mg1 filaments that contain linearmycins. The linearmycins signal intensity is enriched at dense membranous regions, primarily localized to the hyphal branch points. Using fluorescence-lifetime imaging microscopy (FLIM), we determined that the biosynthetic enzymes for linearmycins co-localize to these regions. Thus, we hypothesize that these regions are the active sites of biosynthesis. We used serial FIB-SEM to generate volume reconstructions of *S.* sp. Mg1 wild type and a linearmycin-deficient mutant, revealing subcellular granular structures that are dependent on linearmycin biosynthesis. The linearmycin-deficient mutant strain, which no longer forms the granular structures, displays defects in the cell envelope that are visible using fluorescent dyes and 3D volume reconstructions. The results presented in this study reveal the dependence of cell envelope integrity on the production of linearmycins, which are membrane-disrupting antibiotics when applied to other organisms.

## RESULTS

### Fluorescence detection of linearmycins

Linearmycins belong to the class of polyene metabolites (**Fig S1A**) that includes filipins, amphotericin B, and nystatin, which are potent antifungal antibiotics (28). Due to regions of extensive poly-unsaturation, both amphotericin and nystatin exhibit fluorescence that has been used to probe sterol-rich eukaryotic cell membranes (19,29–31). Given their structural similarity, we hypothesized that the linearmycins would exhibit similar fluorescence (20). We purified linearmycins from the wild-type *S. sp* Mg1 strain as described previously (9). Using excitation wavelength of 360 nm, we acquired the fluorescence emission spectrum for pure linearmycins. The emission spectrum shows a broad peak spanning from 430-550 nm encompassing blue and green regions in the visible spectrum with maximal intensity near 490 nm (**Fig S1B**).

Direct detection of linearmycin fluorescence within whole cells requires overcoming background fluorescence common to hyphae of *Streptomyces*. The measured emission spectrum of the linearmycins is compatible with multiphoton microscopy, as described previously for the related polyketide ECO-02301 (20). Thus, three photon excitation will provide a means for specific detection of linearmycins in whole cells. To determine if we could detect 2P and 3P excitation, we used a 1040 nm laser source to excite fluorescence of linearmycins, extracted from *S.* sp Mg1 culture, and then applied to a microscope slide (**Fig S1C**). Three different bandpass filters were used to detect fluorescence as we described previously (20) . We detected the 3P excitation signal using a 400-480 nm filter (blue). The 2P signal was observed using a 490-560 nm (green) filter and a relatively weak signal with a 570-640 nm (red) emission filter. The red signal we suspected is primarily from background fluorescence of contaminating extracted metabolites, while the green signal combines linearmycins fluorescence and background signal (**Fig S1C**). The relative signal intensities correspond with the expected fluorescence emission maxima from single-photon measurements. For this study, we assign 3P-blue, 2P-green, and 2P-red to fluorescence detected using these emission filters, respectively.

### Detection of linearmycins in S. Mg1 filaments

We next sought to visualize linearmycin fluorescence in whole bacterial cells using 2P and 3P detection. The intrinsic autofluorescence from *Streptomyces* samples is a common problem for whole-cell fluorescence detection and can vary with time in culture (10). We used two-day liquid cultures of wild-type *S.* Mg1, when linearmycin synthesis is detected but has not peaked, to visualize fluorescence from mycelia (9). We detected fluorescence emission using the 1040nm laser and three detection filters: 3P-blue, 2P-green, and 2P-red (**Fig 1A**). To identify the signal(s) specific to linearmycins, we used a linearmycin-deficient mutant strain (Δ*lnyI*) of *S*. Mg1. The Δ*lnyI S*. Mg1 strain lacks the first enzyme in the biosynthetic pathway, an acyltransferase, preventing all biosynthetic activity and production of linearmycins, which was confirmed using mass spectrometry (9). Upon 3P activation, Δ*lnyI* strain had no detectable signal in the 3P channel, while fluorescence was detected from 2P excitation with both 2P-green and 2P-red filters (**Fig 1B**). The result, consistent with the extracted linearmycins data, revealed that the 3P-blue fluorescence we observe is specific for linearmycins. The 2P-green signal is likely a combination of linearmycins and unknown autofluorescence, while the 2P-red signal is solely contributed by background autofluorescence outside the spectrum for linearmycins. Thus, by using the combination of band pass filters we can selectively detect linearmycins. To further confirm that loss of the 3P-blue fluorescence was specific to linearmycins, we genetically complemented the Δ*lnyI S*. sp Mg1 by expressing the *lnyI* gene, inserted at the φ-C31 phage *attB* locus (9). The complemented strain, Δ*lnyI::lnyI* restored linearmycin production and 3P-blue fluorescence in the hyphae (**Fig 1C**). Combined, these results demonstrate that linearmycin-specific fluorescence from 3P excitation is found co-incident with the hyphae and separable from other background fluorescence signals.

**Figure 1:**
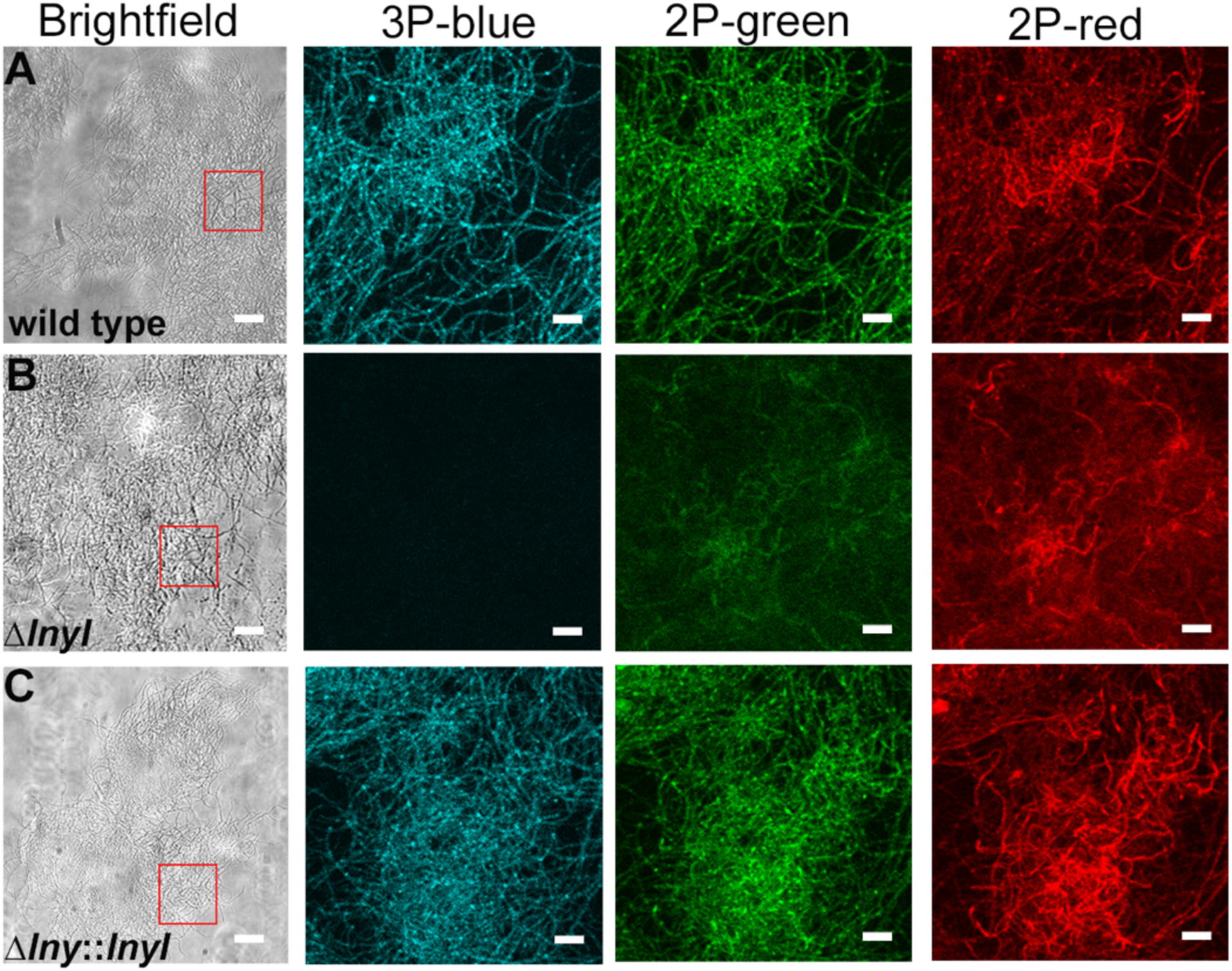
Measurement of 2P and 3P fluorescence in *S.* Mg1 hyphae. Brightfield, 3P excitation detected using 400-480 nm filter (3P-blue), 2P excitation detected using 490-560 nm (2P-green) and 570-640 nm (2P-red) filters for (**A**) wild-type S. Mg1, (**B**) linearmycin deficient Δ*lnyI S*. Mg1, and (**C**) the complement strain Δ*lny::lnyI*. The regions selected from the brightfield image to visualize with 3P/2P fluorescence are depicted with red square. Scale bars for brightfield is 20 μm while the scale bar for fluorescence images is 5 μm.

### Distribution of linearmycins in S. Mg1 filaments and mycelia

Our next objective was to track linearmycins in vegetative hyphae to determine their spatial distribution. Linearmycin BGC expression is first detectable at 24 hours and peaks at 48 hours in liquid culture with linearmyin accumulation following a similar profile (9). Thus, to detect linearmycin signal in live hyphae, we used mycelial masses taken from 48-hour cultures. One advantage of three-photon imaging is the high resolution in biological samples up to millimeter depths, which we observed previously with *S. aizunensis* (20). Using 3P excitation, we could resolve fluorescent signals from individual hyphae within the three-dimensional mycelium. The 2P-red autofluorescence served as a control. With the combined 2P and 3P detection, we identified the linearmycin-specific, 3P-blue fluorescence, at low magnification (2x) in a heterogeneous pattern among the hyphae (**Fig 2A**). Intriguingly, the 2P-red and 3P-blue fluorescence were inversely correlated. Hyphae with intense linearmycin signal exhibited minimal red autofluorescence and vice versa. The difference in signal was most visible using a higher magnification (10x), where we observed within some hyphae that the signals were segregated. Among many hyphae, we observed a pattern of signal separation, which often occured near either cell division planes or branch points for hyphae (**Fig 2B**). These observations suggest that linearmycin biosynthesis is spatially regulated and restricted to a subset of hyphae in the *S*. Mg1 mycelium.

**Figure 2:**
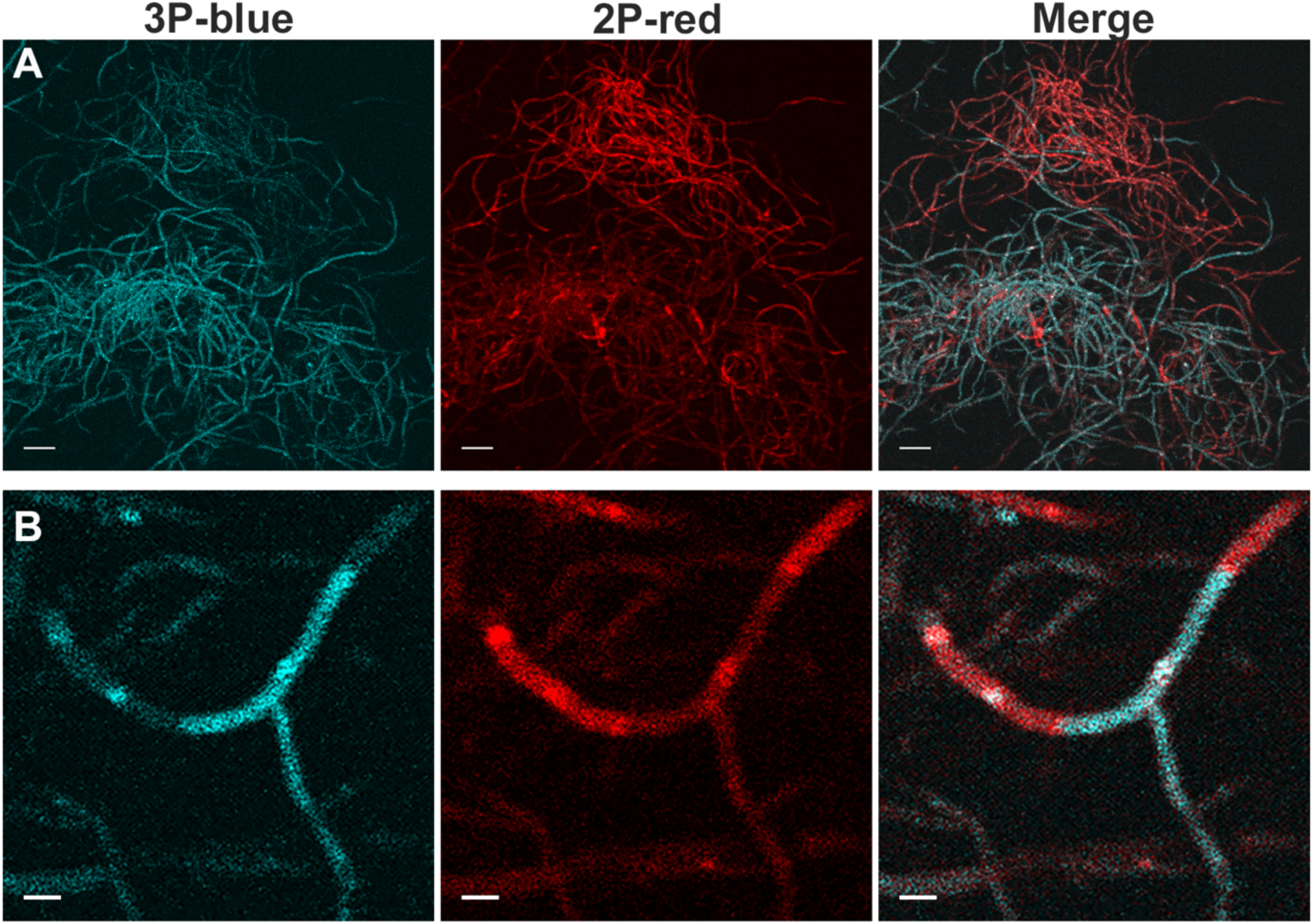
Spatial distribution of linearmycins in *S*. Mg1 mycelium. 3P and 2P excitation fluorescence of *S.* Mg1 hyphae. (**A**) Low magnification (2x) of the mycelium on wet mount, and (**B**) high magnification (10x) of a single hypha from an independent mycelium. 3P-blue represents the fluorescence from linearmycins detected using the blue filter, 2P-red shows the background fluorescence independent of linearmycin fluorescence using the red filter. The images were overlaid and aligned to generate the merged image. Scale bar for low magnification images = 20 μm and scale bar high magnification images = 2 μm.

### Association of linearmycins with membrane-rich regions

3P imaging revealed discrete, high-intensity fluorescence within *S*. Mg1 hyphae, indicating the enrichment of linearmycins in specific regions. Because the linearmycins are lipophilic metabolites, we suspected they may associate with regions of intracellular membrane accumulation. Studies with other *Streptomyces* species demonstrated two types of cross membranes in hyphal cells as detected by fluorescent lipophilic dyes and EM (23). The cross membranes are cytoplasmic extensions of the cellular membrane that either mark the ends of individual cells or appear as non-continuous membrane accumulations within *Streptomyces* hyphae during vegetative growth. We investigated whether the sites of linearmycin accumulation corresponded with either of these patterns by staining the membranes of wild-type *S.* sp Mg1 with the lipophilic dye, FM4-64 (23). FM4-64 fluorescence was detected using the 2P-red filter (570-640 nm), while linearmycins were detected by 3P-blue (400-480 nm) (**Fig 3A**). FM4-64 fluorescence was coincident with cell membranes, and accumulated within intense subcellular foci in a pattern similar to that observed with *S. coelicolor* cross membranes (23). Overlays of the stained 2P-red and linearmycin 3P-blue images do not align perfectly due to hyphal movement in wet mounts during acquisition and filter switching. Nevertheless, the overlaid images provide a visible association of regions with high signal intensity in both stained 2P-red and 3P-blue, indicating that many of the dense membrane accumulations stained by FM4-64 are enriched with linearmycins. Because we observed accumulation of linearmycin signal with the membrane-enriched regions, we selected 5 independent panels for the wild-type strain stained with FM4-64 and manually counted the number of coincident regions of intense linearmycin signal. We counted ∼80% of the total FM4-64-stained regions that coincide with linearmycin-intense regions (**Table S1**).

**Figure 3:**
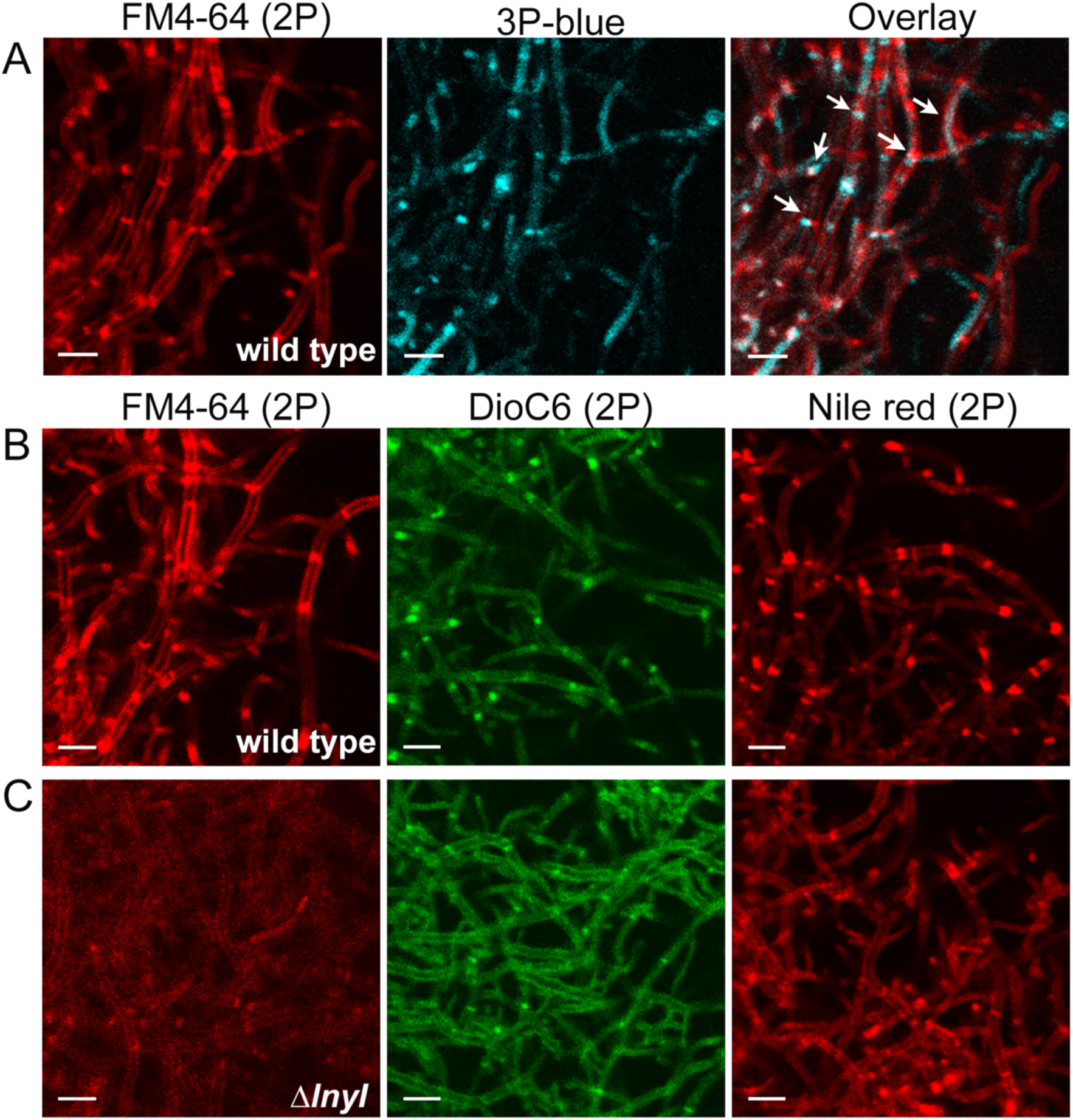
Membrane staining of *S*. Mg1 filaments. (**A**) FM4-64 staining detected using red filter (2P) and linearmycin fluorescence detected simultaneously using 3P-blue filter of the same region in wild-type filaments. Selected regions with colocalized fluorescence are highlighted with white arrows. (**B and C**) Membrane staining of (B) wild type *S*. Mg1 and (C) Δ*lnyI S*. Mg1 filaments using FM4-64, DioC6, and Nile red. (Note: The 3P-blue image of Δ*lnyI, which shows no fluorescent signal,* is available in supplemental information, Fig S2). Binding of FM4-64 and Nile red was detected using the red filter while DioC6 binding was detected using green filter. Membrane-stained cells were imaged using 10% laser power while 3P-blue with 40% laser power. Scale bars = 10 μm.

To show that the 3P-blue fluorescence at membrane-rich regions is linearmycin specific, we stained the Δ*lnyI* mutant strain with FM4-64. Compared to the wild-type control, the Δ*lnyI S*. Mg1 filaments exhibited relatively weak and dispersed FM4-64 staining, and the pattern of dense membrane regions was disrupted (**Fig 3B-C**). As expected, the Δ*lnyI* strain showed no 3P-blue signal (**Fig S2A**). FM4-64 preferentially binds anionic lipids and stains the outer leaflet of cellular membranes (32). To determine if the defect is specific to FM4-64, we selected two additional membrane permeable dyes based on their differences in membrane incorporation. Nile red has been shown to bind preferentially to neutral lipids (33,34). DioC6, is a cationic dye that is sensitive to membrane potential (35). 2P-imaging revealed that these dyes (Nile red: 2P-red, DioC6: 2P-green) stain the wild-type *S.* sp Mg1 cell membranes and intense focal regions (**Fig 3B**). Using the Δ*lnyI S*. Mg1, the Nile red and DioC6 retained staining coincident with the membranes, but the intense foci were greatly diminished, like we observed with FM4-64. We counted regions of intense Nile-red staining and 3P-blue linearmycin signal as done for FM4-64 and found again ∼80% of membrane-enriched regions (**Table S1 and Figure S2B**). Therefore, the lipid-rich regions appear to be highly correlated with sites of linearmycin enrichment. The contrasting membrane staining patterns observed between wild type and Δ*lnyI S.* Mg1 strains suggest that the loss of linearmycins perturbs the cell membranes and disrupts the formation of cytoplasmic membrane-dense regions.

### Co-localization of linearmycins with biosynthetic enzyme machinery using FLIM

The focal fluorescence of linearmycins in regions that are strongly stained by lipophilic dyes suggests membrane accumulation may colocalize to regions of linearmycin biosynthesis. To test the prediction, we designed an in-frame translational fusion of the *lnyI* gene with the *ypet* gene (*lnyI-ypet*) to be expressed in Δ*lnyI S*. Mg1 strain. We confirmed that the *lnyI-ypet* functionally complements the Δ*lnyI* strain using linearmycin-dependent lysis of co-cultured *B. subtilis* (**Fig S3**) (9). We monitored the epifluorescence in wild type, the Δ*lnyI* vector control, and the Δ*lnyI::lnyI-ypet* strain using yellow emission filter (Ex: 500 nm; Em: 540 nm) filter to detect Ypet expression. All three *S*. Mg1 strains had high background fluorescence in the yellow channels (**Fig S4**). A notable difference in the LnyI-Ypet expressing strain was a punctate pattern within the filaments, which is commonly observed for autofluorescence from many *Streptomyces* species (36,37). To differentiate the LnyI-YPet signal from background yellow fluorescence, we visualized the strains using multiphoton fluorescence combined with fluorescence-lifetime imaging (FLIM) (**Fig 4**). Notably, excitation of LnyI-Ypet using the 1040 nm laser did not reveal the punctate fluorescence pattern previously observed, indicating that the foci are auto-fluorescent compounds excited only by shorter wavelengths (**Fig S4**).

**Figure 4:**
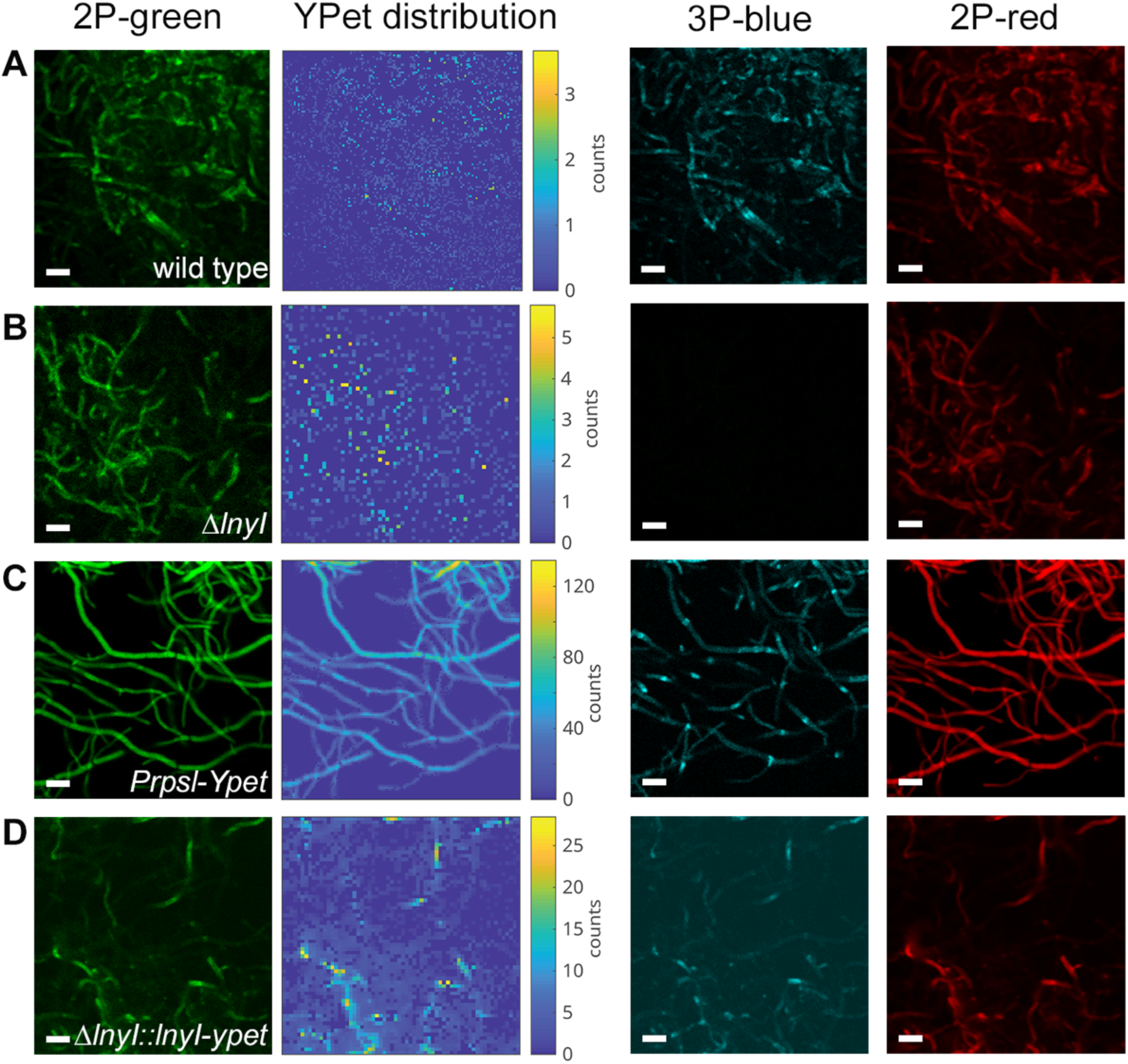
FLIM measurements using various *S*. Mg1 strains. Visualization of fluorescence used to acquire FLIM data with the Time Tagger (Swabian Instruments). Imaging micrographs show fluorescence with green (2P-green), blue (3P-blue) and red (2P-red) filters. Ypet distribution heatmaps reveal exclusively the pixel intensity for the FLIM signal from Ypet (∼3.2 ns). These images were generated for (**A**) wild type, (**B**) Δ*lnyI S*. Mg1, (**C**) the wild-type strain expressing Ypet under a constitutive promoter (P_rpsL_), and (**D**) the Δ*lnyI* strain expressing LnyI-YPet. Scale bars = 5 μm.

Fluorescence lifetime imaging (FLIM) enables differentiation of various fluorophores in the field of view based on fluorescence decay times (38). The data for all the fluorophores in the field of view were collected using a time-to-digital converter that records the relative arrival time of photons at the PMTs across all three emission filters. We generated phasor plots that provide a simplified view of the signal complexity (number of different FLIM lifetimes detected) for each fluorescence channel during the time of exposure (**Fig S5**). Points along the arc indicate single-exponential decay (i.e. single FLIM signal), while deviations from the arc indicate multi-exponential decays, revealing the presence of multiple fluorophores. To record the fluorescence lifetime specific for Ypet in the *S.* Mg1 hyphae, we expressed *ypet* under the strong promoter, P*rpsL*, and monitored the fluorescence using the 2P-green filter (**Fig 4C**). The phasor plot for P*rpsl*-*ypet* was dominated by a uniform signal for the fluorescence lifetime of Ypet (**Fig. S5**). Fitting the data to exponential decay models determined the lifetime of Ypet as ∼3.2 ns (**Table 1**), similar to the reported lifetime of 3.2-3.4 ns (39). Data acquired using the 3P-blue filter were used to generate phasor plots specific to linearmycin fluorescence, which were absent using the *ΔlnyI* strain (**Fig S5**). Decay plots of the FLIM data from wild type revealed three lifetimes: 17 ns, 2.7 ns, and 0.5 ns (**Table 1**). Comparison to the FLIM data from the Δ*lnyI* strain revealed loss of the 17 ns decay signal and the persistence of weak decay lifetimes at 0.5 ns and 2.5 ns. This difference was apparent in the imaging and phasor plots of the Δ*lnyI* FLIM data (**Fig 4, and Fig S5**). Therefore, we recorded the 3P-linearmycin fluorescence lifetime as ∼17 ns. Using the 2P-green filter, we expected signal from the linearmycins along with wild-type background autofluorescence. Under 2P-green detection, we observed three fluorescent lifetimes from the wild-type strain: 14 ns, 2.6 ns, and 0.63 ns (**Table 1)**. We conclude that the 14 +/-2 ns signal arises from linearmycins, because the Δ*lnyI* strain lacks this signal. The complementing strain, Δ*lnyI::lnyI-ypet* restored the 17 ns FLIM signal assigned to linearmycins (**Table 1, Fig S5**). The Ypet signal of 3.2 n was also present in the Δ*lnyI::lnyI-ypet* strain, confirming expression of the *lnyI-ypet* fusion. The phasor plot of the 2P-green signal from the P*rpsL*-*ypet* strain provided a reference for Ypet fluorescence under the experimental conditions used.

**Table. 1.**
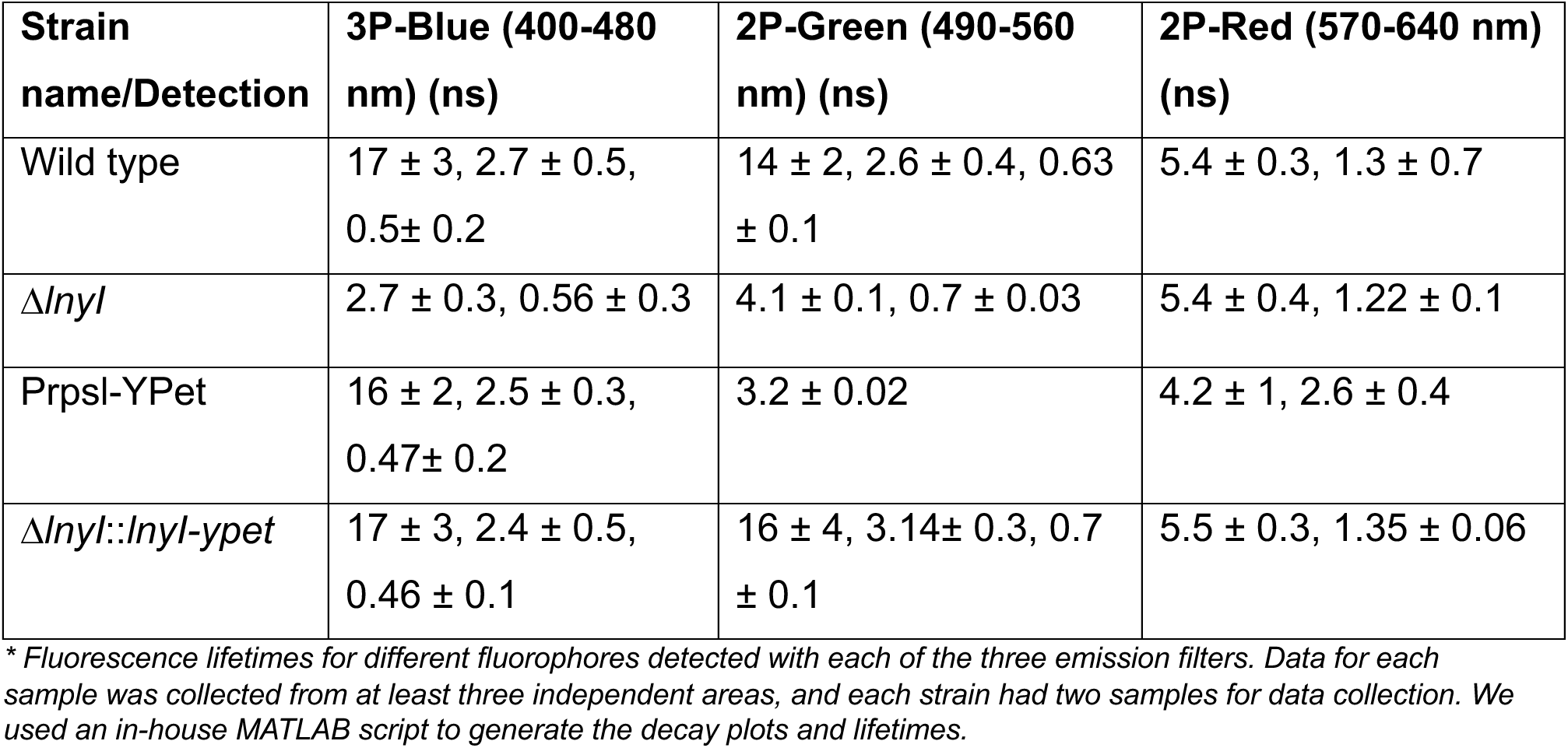
Lifetimes of fluorescent signals detected by FLIM *.

To localize the LnyI-Ypet fluorescence in the complemented strain, we used the 3.2 ns lifetime to generate 2D maps of Ypet fluorescence specifically (**Fig 4**). Wild type and the Δ*lnyI* strains showed only background noise in the 2D maps, as expected due to the lack of Ypet in these strains (**Fig 4A and B**). The distribution of Ypet signal was uniform throughout the filaments in the P*rpsL-ypet* strain as expected for the strong expression from P*rpsL* (**Fig 4C**). Mapping the 3.2 ns signal from the Δ*lnyI::lnyI-ypet* strain, we observed a pattern of Ypet fluorescence in discrete regions of *S.* Mg1 filaments (**Fig 4D**). By comparison, the Ypet fluorescence driven by the *lnyI* native promoter is relatively weak compared to PrpsL-Ypet, as observed in the corresponding phasor plot (**Fig. S5**). Nevertheless, we observed a pattern of relatively intense signal coincident with 3P-blue linearmycins fluorescence and like that from lipophilic dye staining (**Fig 3 and 4**). Using 2P-red signal as a background reference, we observed no significant change in FLIM signals from red background fluorescence except for the PrpsL-Ypet strain, where we suspect that bleed-through from the strong Ypet expression may distort the data detection in the red channel (**Fig 4, S5**). Collectively, the FLIM data reveals the localization of LnyI-Ypet and indicates that the biosynthetic enzymes for linearmycin synthesis accumulate within intracellular compartments enriched in membrane bilayers and linearmycins.

### 3D-cell reconstructions from volume EM data reveal linearmycin-dependent subcellular compartments and cell envelope defects from loss of linearmycin synthesis

Given that our fluorescence microscopy suggests that linearmycin biosynthesis is compartmentalized to membrane-rich subcellular regions, we sought a high-magnification approach to identify relevant structural features. Because imaging the filamentous mycelia at two days of growth is impractical using standard thin-section TEM, we used volume EM to generate high-resolution 3D reconstructions of *S.* Mg1 mycelia. Cultures of wild type, Δ*lnyI*, and Δ*lnyI::lnyI* complemented strains were chemically fixed, stained with uranyl acetate and OsO4, and embedded in thin layer of resin (40). We used Focused Ion Beam-SEM (FIB-SEM) and generated volume imaging data sets of mycelia that were iteratively milled and imaged several hundred times **(Figs S6 and S7)**. Individual SEM images from the data set revealed some notable differences between the cell envelopes of wild type and Δ*lnyI S*. Mg1. For example, the wild-type cell envelope has higher intensity and a more uniform thickness than that of the Δ*lnyI* mutant strain (**Fig S8**). One advantage of serial FIB-SEM imaging is the ability to generate 3D volumes from alignment and compiling of the individual SEM images (**Fig S9**). From these digital data sets, we selected subsets of images to capture individual hyphae for each of the strains (**Fig S10**). We then segmented the cell envelope and subcellular features based on their relative signal intensity. Image sets for each *S.* Mg1 strain were processed identically to provide comparable 3D volumes of cells and subcellular features (**Fig 5 and Fig S11**). Reconstructions of hyphae revealed distortions on the cell surface of the Δ*lnyI* mutant strain, whereas the wild type and the Δ*lnyI*::*lnyI* complementation strain had smooth surfaces (**Fig 5A-C, Fig S11**). Although we noted some differences in the cell envelopes when comparing cells from single SEM images, the extent of the surface distortions was apparent only in the volume reconstructions. To observe subcellular features of the strains, we rendered the cell envelope transparent or, alternatively, removed a section of the cell envelope from the 3D digital reconstructions (**Fig 5D-F, Fig S11**). Prominent among many of the wild-type cells were globular densities that were absent in the Δ*lnyI* mutant strain. The subcellular structures appeared to coalesce in discrete regions, often observed near hyphal branch points. Although generating a statistical correlation using the EM data is impractical, the pattern observed is consistent with that observed with membrane staining and 3P-blue linearmycin signal (**Fig 5A, Table S1**). In contrast, we never observed these subcellular densities in the identically processed Δ*lnyI S*. Mg1 hyphae (**Fig 5E, Fig S11B**). However, they were restored in the Δ*lnyI*::*lnyI* complemented strain, demonstrating their dependence upon functional linearmycin biosynthesis **(Fig 5F and Fig S11C)**. Thus, we concluded that the densities observed are linearmycin-dependent structures. The observed cell-surface defects are a phenotype of *lnyI* deletion, exhibited in the absence of linearmycins, and appear consistent with changes in membrane staining observed using fluorescence. The observation of linearmycin-dependent subcellular structures, and the defects associated with loss of linearmycin biosynthesis, suggests that linearmycins, in addition to their antibiotic function, are an integral component of the bacterial cell envelope in this organism.

**Figure 5:**
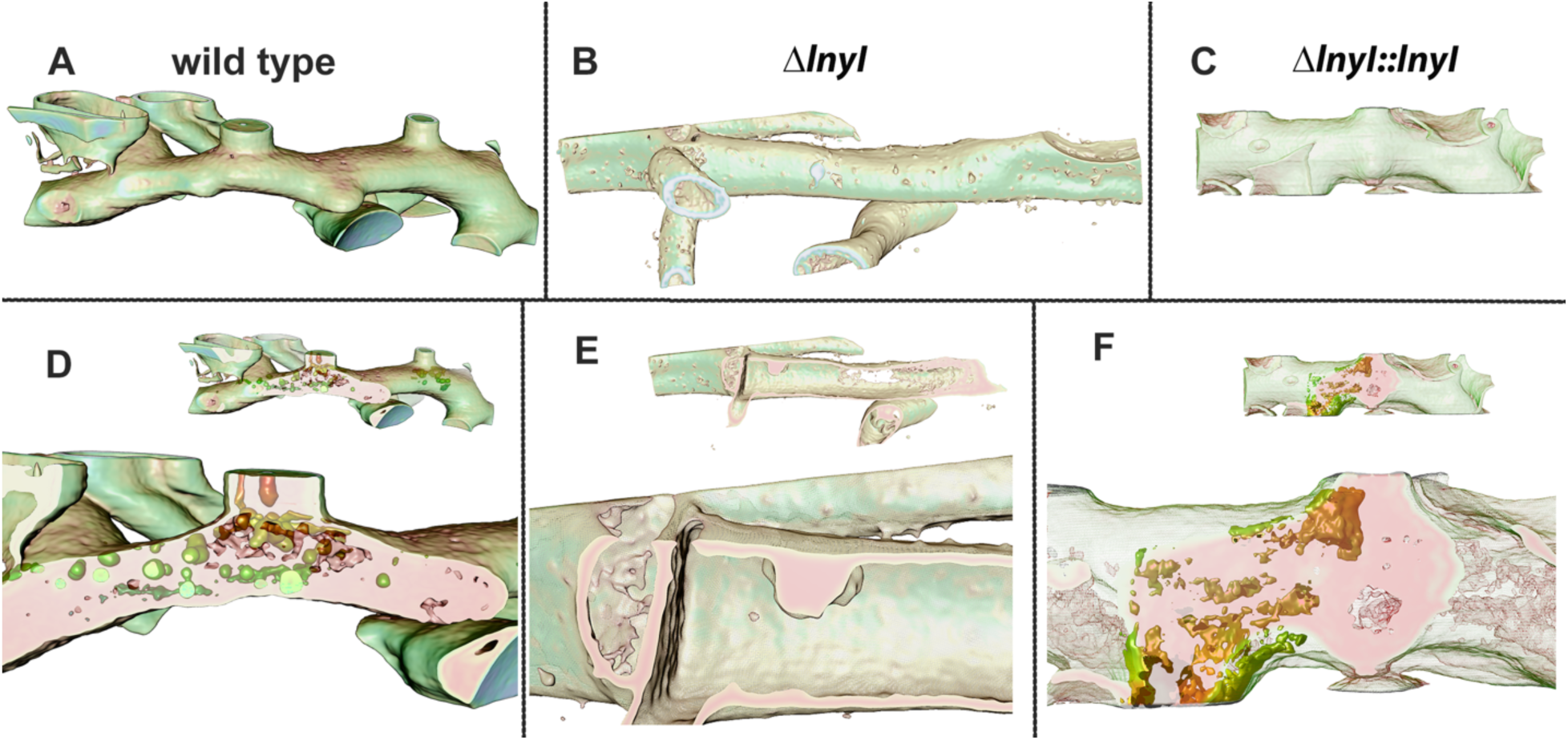
Volume FIB-SEM reconstructions of *S*. Mg1 filaments. Three-dimensional reconstructions of individual filaments from the image datasets collected using FIB-SEM (voxel size 5 x 5 x 10 nm). Segments of individual hyphae were separated in silico from proximal hyphae and hyphal branches and rendered to visualize the cell surfaces of (**A**) wild type, (**B**) the Δ*lnyI* strain, and (**C**) the Δ*lnyI*::*lnyI* complemented strain. Panels D-F show the same hyphal segments with a longitudinal section of the cell surface removed to reveal the underlying subcellular contents. Panels are (**D**) wild type, (**E**) the Δ*lnyI* mutant strain, and (**F**) the Δ*lnyI*::*lnyI* complemented strain. Each panel has a small inset of the same segment from A-C for reference. Enlarged images are shown below to reveal granular subcellular structures that are present in wild type and the complement strain, but absent in the Δ*lnyI* mutant strain. Color parameters applied identically during processing reveal differences in EM signal intensity. The diameter of individual hyphae varies between cells and branches with a range ∼0.5-1.5 µm. For reference, see Fig S10 with scale bars from individual micrographs. Movie files are included in the supplemental materials to provide multiple view angles for each hyphal segment (Movies S1-S3).

## DISCUSSION

Streptomycetes produce polyene antibacterial and antifungal agents, including the linearmycins, amphotericin B, and nystatin (41,42).These polyene antibiotics target the cell membranes of exposed organisms and have been used extensively as chromophores to visualize drug-membrane interactions (17–19). The linearmycins, amphotericin B, and other *Streptomyces* metabolites are encapsulated within extracellular vesicles of their producing organisms (9,43–46). A key finding for our current work was the earlier observation that a genetic disruption of linearmycin biosynthesis disrupts extracellular vesicle formation, indicating a direct link between the two processes (9). To explore the connection between linearmycins and the *Streptomyces* cell membrane, we developed a method to visualize fluorescent metabolites within mycelia (20). Using two- and three-photon excitation fluorescence microscopy, we can image mycelia with single-cell resolution up to a depth of 750 μm (20). Focusing on *S.* Mg1, we found that linearmycin fluorescence is specifically detectable with three-photon excitation (400–480 nm), while other fluorophores are detected with two-photon excitation (**Fig 1**). We used linearmycin fluorescence in combination with lipophilic dyes and a fluorescent protein to track the biosynthesis and distribution of the metabolites in whole cells (**Fig 2, 3 and 4**).

We observed a heterogenous distribution of linearmycins among bacterial hyphae. Linearmycin fluorescence was found in a subset of hyphal cells within the mycelia, with intense accumulation of signal in localized regions, notably at the branch points of the filaments (**Fig 2**). Lipophilic dyes accumulated in these regions, consistent with localization of intracellular membrane structures (23). Recent studies on *S. venezuelae* showed that flavin-containing proteins contribute much of the green autofluorescent foci observed in the vegetative mycelia (36,37). We therefore suspect the intense, punctate yellow foci observed in *S*. Mg1 vegetative filaments may be due to flavin autofluorescence. However, FLIM data mapped to images revealed that the LnyI-Ypet fusion protein frequently co-localizes with the regions of intense linearmycin fluorescence, suggesting that a subset of these regions are biosynthetic centers for production of linearmycins and their incorporation into cellular membrane (**Fig 4**). Our results have broad ramifications for microscopy of filamentous microorganisms, and complement parallel approaches such as Raman microscopy, previously used to detect amphotericin B (15). An important advantage of 3P-fluorescence imaging is the minimal noise from light scatter and overlapping fluorescence detection. Related metabolites to linearmycins, nystatin and filipin, are used as probes for yeast membranes (31) and cholesterol-rich regions of eukaryotic cells (19), which could be enhanced by 3P excitation to improve signal to noise.

Genetic disruption of linearmycin biosynthesis, which blocks EV biogenesis, produced changes in cell envelope that we detected with fluorescence microscopy and volume EM (**Fig 3 and 5**). The mutant strain, Δ*lnyI*, showed reproducible defects in staining of cell membranes, which we suspect result from a lack of dye incorporation, due to perturbation of wild-type membrane composition. Although the difference could arise due to chemical or redox changes in the mutant cell membranes (26,32). All three lipophilic dyes used revealed a substantial depletion of intense regions of staining (**Fig 3**). This result is consistent with the formation of these regions being dependent upon linearmycin biosynthesis in *S*. Mg1. Not all species of *Streptomyces* produce linearmycins or similar polyene metabolites. The observed dependence of the intensely stained regions on linearmycins suggests that they may not be equivalent to intracellular membranes observed in non-linearmycin producing species (23). However, this would not preclude the linearmycin-dependent structures from associating with other membranous structures in the cells. Thus, we investigated the cellular architecture of wild-type and linearmycin-deficent cells to provide a complementary, high-resolution means of assessing differences between strains.

Using serial FIB-SEM imaging, we generated 3D volume reconstructions of mycelia, and segmented features of differing density within cell filaments. The data enabled us to identify regions where granular compartments accumulated, typically near branch points. These compartments were dependent upon linearmycin synthesis and were not present in the Δ*lnyI* mutant strain (**Fig 5**). Additionally, the mutant strain had distortions in the cellular surface, which appeared as blebs along the surface of the reconstructed volumes. The surface distortions were not present in wild type or the Δ*lnyI::lnyI* complemented strain. The fluorescence and volume EM data indicate that the linearmycins are functional components of membranes within subsets of hyphal cells that produce them. The incorporation of linearmycins into membranes is particularly intriguing because they are membrane disrupting, causing lysis of both *Bacillus subtilis* and unilamellar vesicles in vitro (45).

The results of this study suggest that linearmycins are involved in the maintenance of the membrane integrity of *S.* Mg1 filaments, and that their absence leads to alterations in membrane properties and stability. Membrane biogenesis is a dynamic process, responsive to changing demands of metabolism and environment. For example, a study of *S. coelicolor* showed that membrane lipid composition varies in response to changes in media composition, demonstrating how these bacteria can adapt and modify membrane composition based on metabolic demands (47). Biosynthesis of linearmycins draws heavily on pools of malonyl-CoA and shares this demand with lipid biosynthesis. Increased levels of malonyl-CoA alter lipid metabolism and membrane composition in other organisms (48). Thus, a reasonable explanation for the linearmycin phenotypes observed is that the absence of linearmycins destabilizes overall lipid metabolism, thereby disrupting membrane integrity through bulk changes in lipid composition and membrane integrity. We speculate that the coordinated control of membrane biogenesis and secondary metabolism of linearmycins is fundamental to the control of extracellular vesicle biogenesis, although this model will require additional study.

In summary, we present the first report of direct detection of cellular linearmycins in *Streptomyces* hyphae via 3P-fluorescence microscopy. Further, we report the volume rendition of *Streptomyces* filaments to identify subcellular compartments that require linearmycin biosynthesis and that we hypothesize may be regions for biogenesis of extracellular vesicles. These volume renditions allowed us to detect subcellular structures at nano scale, which has not been attempted with *Streptomyces* earlier. Our findings link linearmycin production to membrane structure and integrity, suggesting that these specialized metabolites not only provide competitive advantages but are also have an integral role in cell physiology and morphology of the producing bacteria.

## Methods

### Bacterial strains and chemical reagents

All the strains used in the study are described in Table S2. *Bacillus subtilis* and *E. coli* strains were cultured in LB (1% w/v tryptone, 0.5% yeast extract and 0.5% NaCl). *Streptomyces* strains were cultured using MYM media (0.4% w/v D-maltose, 0.4% w/v yeast extract, 0.4% w/v malt extract) buffered with 100 mM MOPS pH 7. AS1 plates (0.1% w/v yeast extract, 0.02% L-alanine, 0.02% L-arginine free base, 0.05% L-asparagine, 0.5% w/v soluble starch, 0.25% NaCl, 1% Na_2_SO_4_, 10 mM MgCl_2_, 1.5% Agar) were used for *S*. Mg1 and *E. coli* ET12567 conjugations. Apramycin, chloramphenicol, hygromycin antibiotics were used at standard concentrations for *E. coli* and *S.* Mg1. Δ*lnyI* and PrpsL-YPet *S*. Mg1 strains were grown and sporulated in the media containing apramycin (50 μg/mL). Δ*lnyI::lnyI* and Δ*lnyI::lnyI -ypet S*. Mg1 strains were grown and sporulated in the media containing apramycin (50 μg/mL) and hygromycin (50 μg/mL). All the strains were grown in the absence of any antibiotic for all the microscopy experiments.

### Extraction of linearmycins

Isolation and purification of linearmycins from *S*. Mg1 was carried out using the procedure described previously (9). 1 L buffered MYM7 media containing Diaion HP-20 resin (20 g) was inoculated with *S*. Mg1 spores (10^7^ spores /mL) and incubated for 48 h at 225 rpm and 30°C. The resin was washed and eluted with methanol and dried under vacuum. The dried methanol extract was resuspended in 50% ACN and loaded onto a C8 SPE cartridge (Waters Corp, USA). Methanol gradient from 40%-90% was used to carry out elution and the fractions collected were tested for lysis activity against *B. subtilis*. 90% methanol fraction was active and was used for HPLC separation on Vanquish core (Thermo Fisher Scientific, USA). The sample was loaded onto Phenomenex Luna C18 column and chromatography was carried out using 10 mM ammonium acetate (pH 5) and acetonitrile. The acetonitrile gradient used was 40% for 5 min, 50% for 10 min, 75% for 5 min and back to 40% at 20 min. Elution of linearmycins was monitored at 330 nm. Linearmycin elution was observed around 13 min and 16 min corresponding to A and B forms. Those fractions were pooled, dried, and tested for bioactivity.

### Construction of 11lnyI::lnyI -ypet and PrpsL-Ypet S. Mg1 strains

The plasmid to construction the *lnyI-ypet* fusion was constructed using Gibson assembly with 3 linear fragments. The primers used in the generation of the plasmid are listed in Table S3. Fragment-1 was amplified from pOOB4O7 using primers pOOB40-FP and pOOB40-RP. Amplification of *lnyI* along with its promoter used primers *lnyI*-FP and *lnyI*-RP (fragment-2). A linker sequence was added at the end of the gene to link Ypet to LnyI upon translation. Fragment-3 was obtained upon amplification of the *ypet* gene from pUC17-YPet using the primers *lnyI-yfp*-FP and pOOB40-*yfp*-RP. The concentration for each of the fragments was determined on Nanodrop. The three fragments were stitched together using Gibson assembly (NEB, USA). The reaction mix was transformed into *E. coli* DH5α and plated on LB media containing hygromycin (50 μg/mL). PrpsL-Ypet construct was generated using Gibson assembly with 3 linear fragments. pSET152 vector was amplified using the primers BCH0067 and BCH0068 to generate linear fragment. Amplification of *ypet* and promoter of *rpsL* was carried out using primer pairs BCH0070/0071 and BCH0072/0073 respectively. The three linear fragments were ligated using Gibson assembly (NEB, USA). The reaction mix was transformed into *E. coli* DH5α and plated on LB media containing apramycin (30 μg/mL). Plasmids were isolated from positive colonies and confirmed using DNA sequencing (Eton BioSciences, CA, USA). The confirmed plasmids were transformed into conjugation efficient *E. coli* ET12567 cells and then into Δ*lnyI* and wild type *S*. Mg1 strains through conjugation. Exconjugants were selected on media containing appropriate antibiotics.

### Multiphoton microscopy

*S*. Mg1 strains were grown in 25 mL MYM7 buffered media for the required amount of time. 1 mL of the culture was harvested and resuspended in 150 μL water. 3 μL of the sample was added to the microscope slide and secured with a coverslip. To protect the sample from dehydration, the edges of the coverslip were sealed with vacuum grease, and the slide was used for multiphoton microscopy.

The setup used for multiphoton imaging is based on the open frame design of the modular in-vivo multiphoton microscopy system (MIMMS, https://www.janelia.org/open-science/mimms-10-2016) and described in detail earlier (20,49). In this work we used the high-NA microscope objective (Nikon MRD71970, 100×, 1.45 NA oil) that was used by Classen et al. Blue (ET440/80m-2p, Chroma), green (ET525/70m-2p, Chroma) and red (ET605/70m-2p, Chroma) emission filters are used to select fluorescence emitted in three wavelength ranges (400-480 nm, 490-560 nm and 570-640 nm, respectively). The fluorescence signal is digitized using a fast analog-to-digital converter (ADC) that allows detection of high dynamic range signals synchronized with the laser pulses. Z stack images were acquired for every 0.25 μm and at an average of 25 frames per slice in two channels at a time. Raw images were processed and normalized using ImageJ (FIJI, USA) image editing software.

### Multiphoton Fluorescence Lifetime Imaging (FLIM) measurements of S. Mg1 strains

For the FLIM measurements, the above fluorescence signal is digitized using a time-to-digital analyzer that allows for count rates up to 90 Mtags/s (TimeTagger Ultra, Swabian Instruments) with an RMS timing jitter of 42 ps. This allows to record fluorescence decay curves for each detection channel simultaneous for each pixel. Custom MATLAB scripts are used to convert the TimeTagger data into images, and to analyze the FLIM data. Lifetime data for wild type, PrpsL-Ypet and Δ*lnyI*::*lnyI-ypet* were analyzed using *bin4,* while lifetime data for Δ*lnyI* was analyzed *bin8* due to low signal intensity.

### Chemical fixation and resin embedding of S. Mg1 strains

Spores of *S.* Mg1 strains (10^7^ spores/mL) were inoculated in 25 mL of buffered MYM7 and incubated at 30 °C at 225 rpm for 48 h. 2 mL cells were harvested and washed twice with 1mM MOPS buffer (pH 7.0). All the steps including fixation, staining, dehydration and resin infiltration were carried out using microwave-based protocols to improve efficiency and reduce the time for the treatments (50). The pellet was treated with 500 μL of 2% glutaraldehyde in 0.1 M sodium cacodylate buffer for chemical fixation. The chemically fixed pellet was treated with 1% (w/v) OsO_4_ with 1.5% potassium hexacyanoferrate (II) in 10 mM MOPS buffer (pH 7) followed by 5 washes with water and then the pellet was stained with 1.5% uranyl acetate followed by washes with water (6 times). The stained pellet was dehydrated with 30%, 50%, 70% and 100% ethanol. The step with 100% ethanol was repeated twice. The pellet samples were infiltrated with Spurr’s resin mixture (ERL 4206-5g, DER-3g, NSA-13g and BDMA – 0.2g) in the ratio 1:2, 1:1, 2:1 with ethanol and finally twice with 100% resin (40).

To attain embedding in thin layer of resin, we transferred the resin-infiltrated sample onto 0.45 μm nylon membrane. A few drops of 100% resin were added to the sample. The membrane with the sample was subjected to a low vacuum to slowly remove the resin through the membrane until a thin layer of resin is retained around the sample. The membrane is then left for resin polymerization at 60 °C for 2 days. The polymerized embedded samples were used for FIBSEM image acquisition.

Two-day old bacterial cells were chemically fixed using glutaraldehyde followed by treatment with OsO_4_ and staining with uranyl acetate. A thin layer of Spurr’s resin was coated on the dehydrated cells as mentioned in the Methods section. Upon polymerization of the resin, the samples were sent to Thermo Fischer Scientific imaging facility at Nanoport Hillsboro, OR, for image acquisition using Scios2. An area of interest (AOI) was selected and a thin protective layer of platinum deposited on the sample. Initially, the AOI was milled with low voltage FIB beam (**Fig S6**).

### Focused Ion Beam Scanning Electron Microscopy (FIBSEM) of S. Mg1 strains

Resin embedded *S.* Mg1 strains were sent to Thermo Fisher’s Electron Microscopy Facility Hillsboro NanoPort for FIBSEM data collection. The parameters used for FIBSEM data collection are listed in Table S4.

### Image processing and three-dimensional reconstruction of S. Mg1 filaments

Stacks of micrographs for a given data set were aligned, using Amira and Avizo (ThermoFisher) and ImageJ (FIJI). The aligned stacks of micrographs were then visualized in “bshow” of the software package BSOFT (51). The slices which did not show good alignment were aligned with their predecessor and successor images using “bseries” of BSOFT package.

Regions of interest in the aligned stacks of micrographs were then cropped off and subjected to realignments using the methods mentioned earlier, which gave better results. Segmentations of the regions of interest were then conducted by selecting appropriate threshold range and employing a watershed algorithm implemented in “bflood” of BSOFT package. Software packages UCSF Chimera (52), Avizo and some home written scripts were also used to carry out segmentation. The density maps were subjected to band-pass filtering to remove the high frequency noise signals using “bfilter” (b). The segmented density maps provided extensive particularities about the connections and the interlinks between the subcellular moieties. Visualization of the maps was done in UCSF Chimera.

## Author Contributions

The study was conceptualized by PDS and KJ. The experiments and analysis were carried out by KJ, AC, AV, AF and AS. Manuscript is written by KJ and PDS with excerpts by AV and AS.

## Acknowledgements

We thank the technical support at ThermoFisher Nanoport facility, Hillsboro, OR, USA for FIBSEM data acquisition and Ovation Datasciences, Houston, for providing the data storage and processing platform to generate volume renditions. We are thankful to Dr. Stanislav Vitha at the Microscopy and Imaging Center, TAMU for his help in FIBSEM sample preparation. We acknowledge the contribution of the undergraduate trainees in the lab, Anthony Rudd and Lauren Skrobarczyk, in generation of the plasmids and conjugation into *Streptomyces sp*. Mg1. Research reported in this publication was supported by the National Institute of General Medical Sciences of the National Institutes of Health under award number R01GM141700.

