## supplementary figures 1-11 and tables 1-4 for "Direct fluorescence detection and volume electron microscopy reveal a role for antibiotic biosynthesis in the bacterial cell envelope of *Streptomyces sp.* Mg1"

**Supplemental information: Figures S1-S11, Tables S1-S4, Movies M1-M3**

### Table of Contents

*Figure S1*

*Figure S2*

*Figure S3*

*Figure S4*

*Figure S5*

*Figure S6*

*Figure S7*

*Figure S8*

*Figure S9*

*Figure S10*

*FigureS11*

*Table S1*

*Table S2*

*Table S3*

*Table S4*

*Movie M1*

*Movie M2*

*Movie M3*

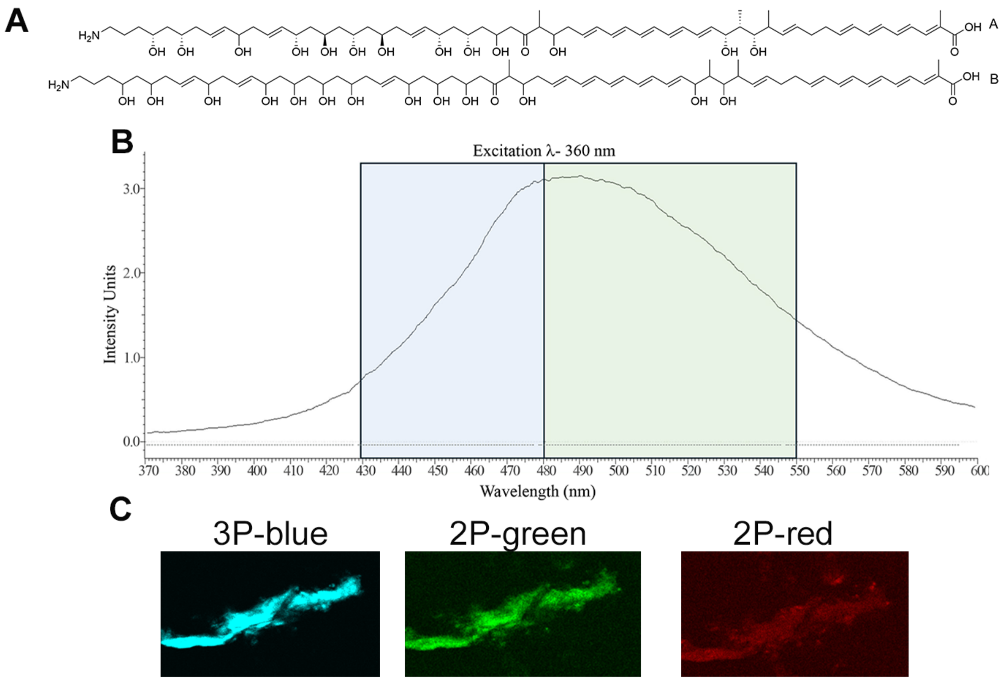


**Figure S1: Linearmycins fluorescence spectrum.** (**A**) Chemical structure of linearmycins A and B. (**B**) Fluorescence emission spectrum measured from extracted linearmycins using an excitation wavelength = 360 nm. The shaded areas show the emission range in the blue and green spectrum respectively. (**C**) Simultaneous 3P-2P excitation fluorescence of extracted linearmycins. 3P-blue depicts the emission detected with the blue filter upon three photon activation, while signal from two-photon excitation is shown using 2P-green and 2P-red filters.

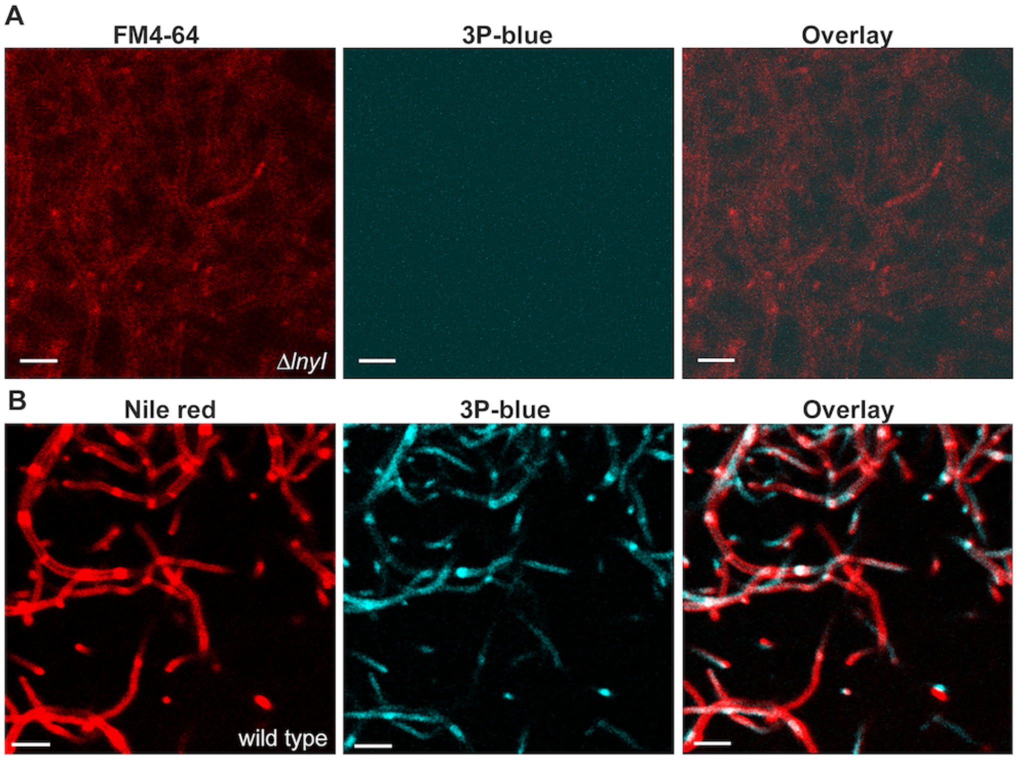


**Figure S2: Membrane staining of S. Mg1 strains.** (**A**) FM4-64-stained Δ*lnyl* hyphae detected with red filter and linearmycin signal with 3P-blue. Linearmycin signal is absent in the Δ*lnyl* strain. (**B**) Nile red staining of wild type *S*. Mg1 hyphae detected as 2P-red and linearmycins signal detected with 3P-blue. The overlay image shows the overlap of the intensely stained membranous regions with linearmycin-intense regions. (Scale bars = 10 μm).

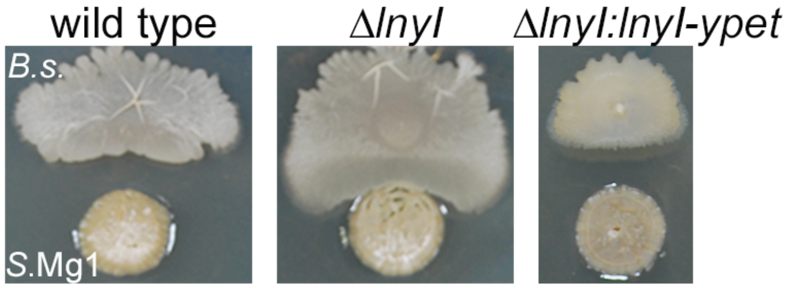


**Figure S3: Bioassay with B. subtilis**. Lysis assay with *B. subtilis* was carried out with wild type, Δ*lnyI* and the Δ*lnyI::lnyI-ypet* complemented strain. Exponentially growing *B. subtili*s cells were spotted 24 hours after spores of S. Mg1 were inoculated. The void area between colonies, viewed after 24 hours co-incubation, is caused specifically by linearmycins, revealing the functional complementation using LnyI-Ypet. *B. subtilis* (*B.s.*) and *S.* Mg1 colonies are labeled, respectively.

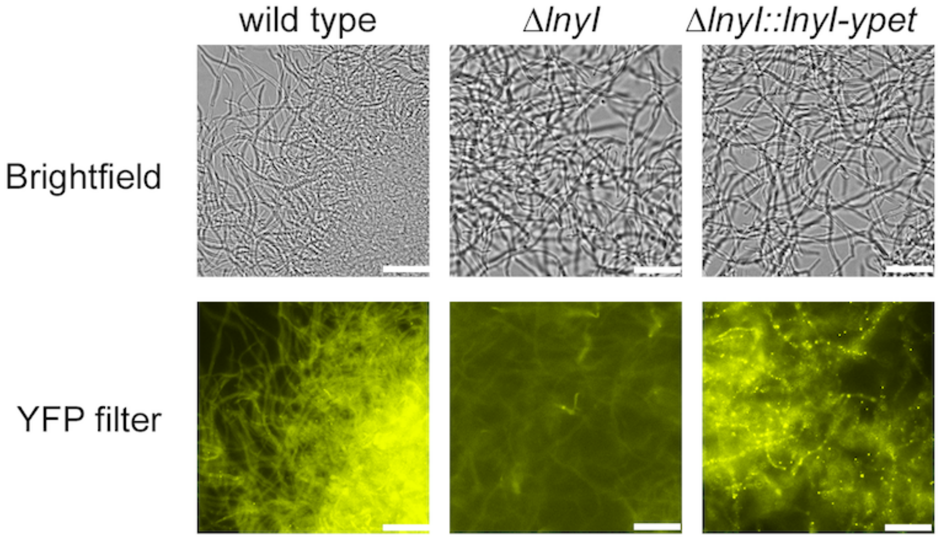


**Figure S4: Epifluorescence in *S*. Mg1 strains.** Epifluorescence detected using YFP filter (Ex 500 nm/Em 542 nm) in two-day old cultures of wild type, Δ*lnyI* and Δ*lnyI::lnyI-ypet S*. Mg1 strains. The images were acquired using an Agilent Lionheart microscope. Scale bar = 10 μm.

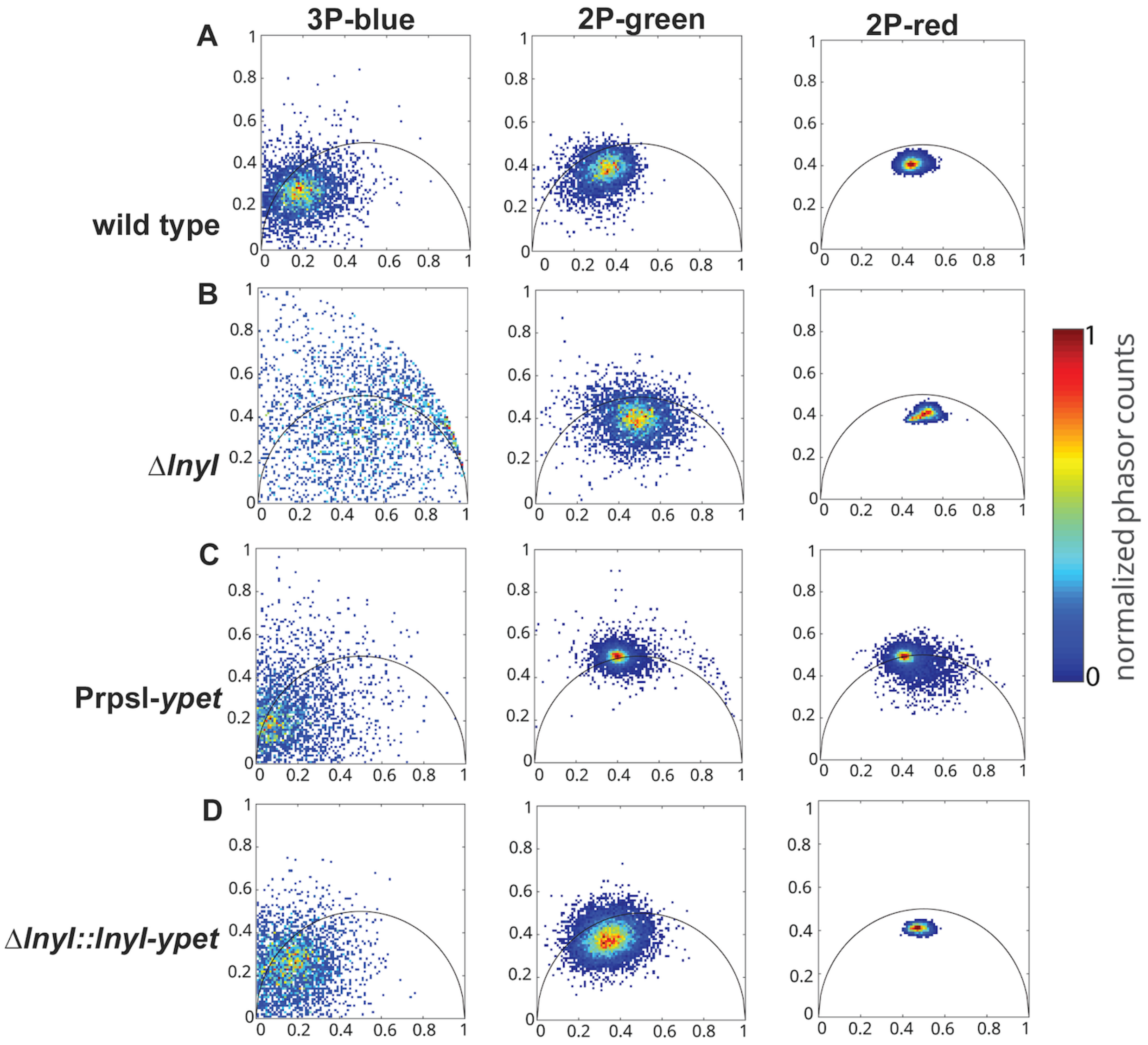


**Figure S5: Phasor plots depicting the FLIM data**. Phasor plots of fluorescence lifetime distributions with 3P-2P excitation obtained from all three emission filters (3P-blue, 2P-green and 2P-red) for (A) wildtype (B) Δ*lnyI S*. Mg1 (C) PrpsL-Ypet and (D) Δ*lnyI* expressing LnyI-Ypet (Δ*lnyI*::*lnyI-ypet*) *S*. Mg1 hyphae. Each point represents the coordinates of a pixel. PrpsL-Ypet was used to identify the Ypet fluorescence lifetime. Lifetime data for wild type, PrpsL-Ypet and Δ*lnyI*::*lnyI-ypet* were analyzed using *bin4* while Lifetime data for Δ*lnyI* was analyzed *bin8 to compensate for low signal intensity*.

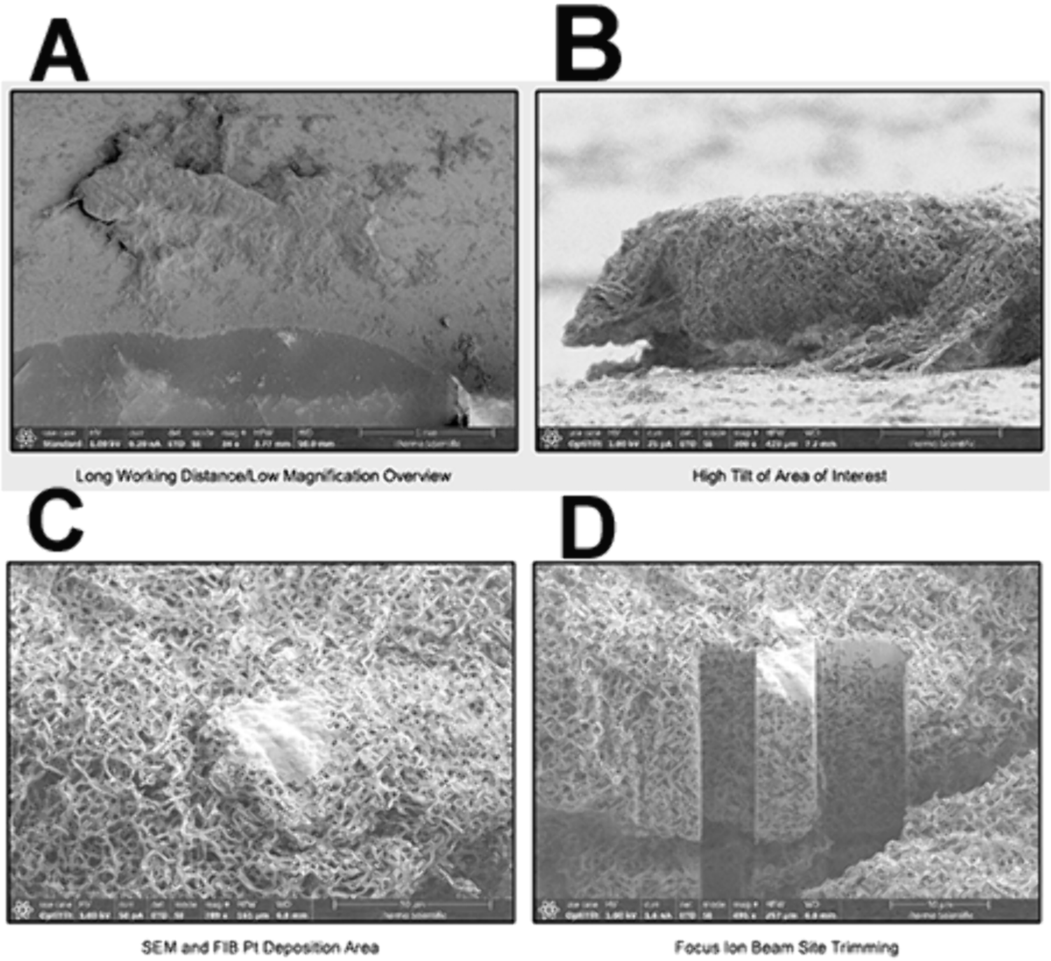


**Figure S6: Sample processing for FIBSEM data acquisition. (**A) Wild type *S*. Mg1 sample embedded in thin layer of Spurr's resin. (B) High tilt view of the region of interest. (C) SEM of the region for FIB milling after Pt deposition. (D) FIB milling at the region of interest.

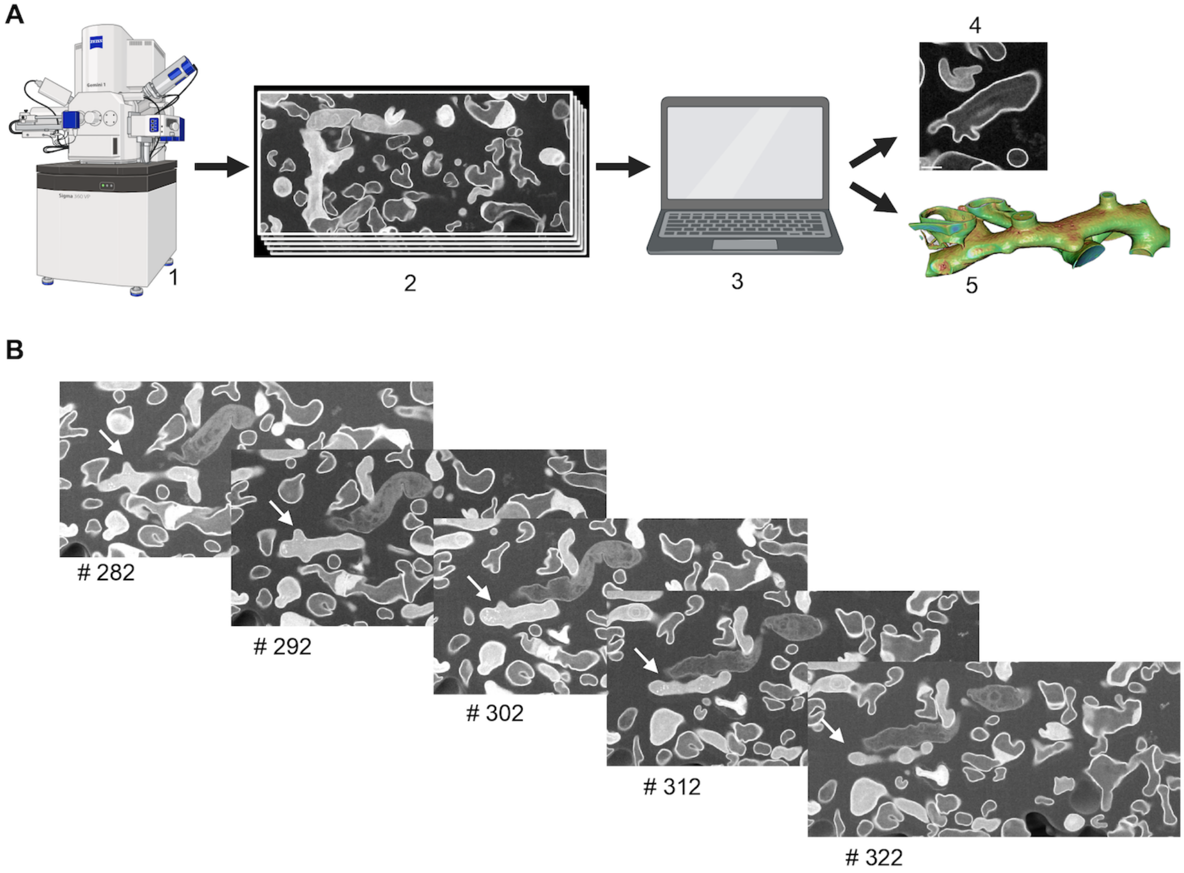


**Figure S7: Schematic of data acquisition and processing of FIBSEM datasets.** (A) The serial FIB-SEM workflow for imaging of embedded samples. (1) FIB milling and SEM generates (2) a series of images at 7-10 nm step size. The images are then (3) computationally aligned and processed to generate 3D-volumes. The data may be viewed as (4) individual images or segmented and processed to visualize (5) 3D reconstructions of individual hyphae. (B) The bottom panels show representative slices from an image dataset for the wild-type strain, showing the continuity of the selected hypha (white arrow) used for 3D reconstruction.

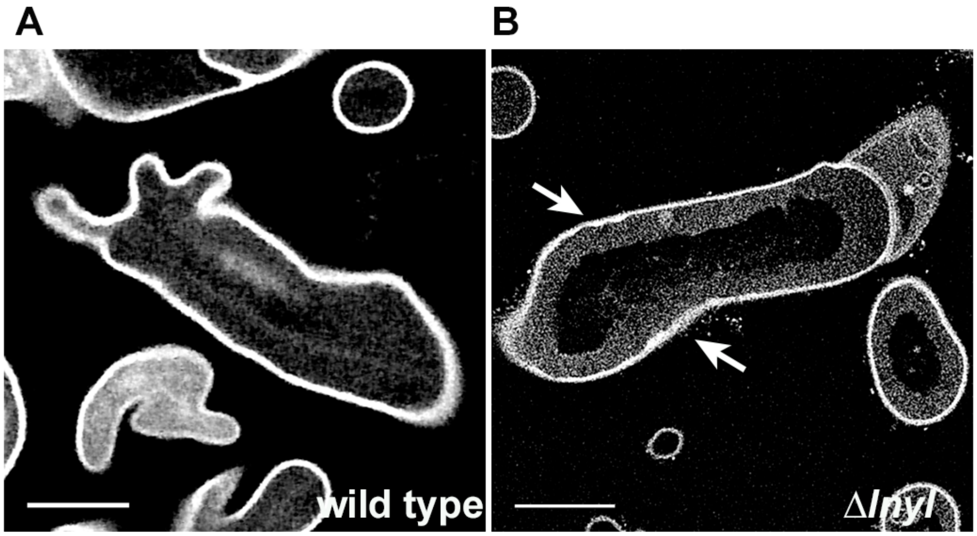


**Figure S8**: **Comparison of filaments from wild-type and Δ*lnyl S*. Mg1 strains**. Back scatter detector (BSD) images for (**A**) wild type and (**B**) the Δ*lnyl S*. Mg1 strain were normalized for contrast to compare the cell envelopes in both the strains. Arrows highlight examples of areas in the mutant strain with abnormal cell envelope uniformity and thickness. Scale bar = 1 μm.

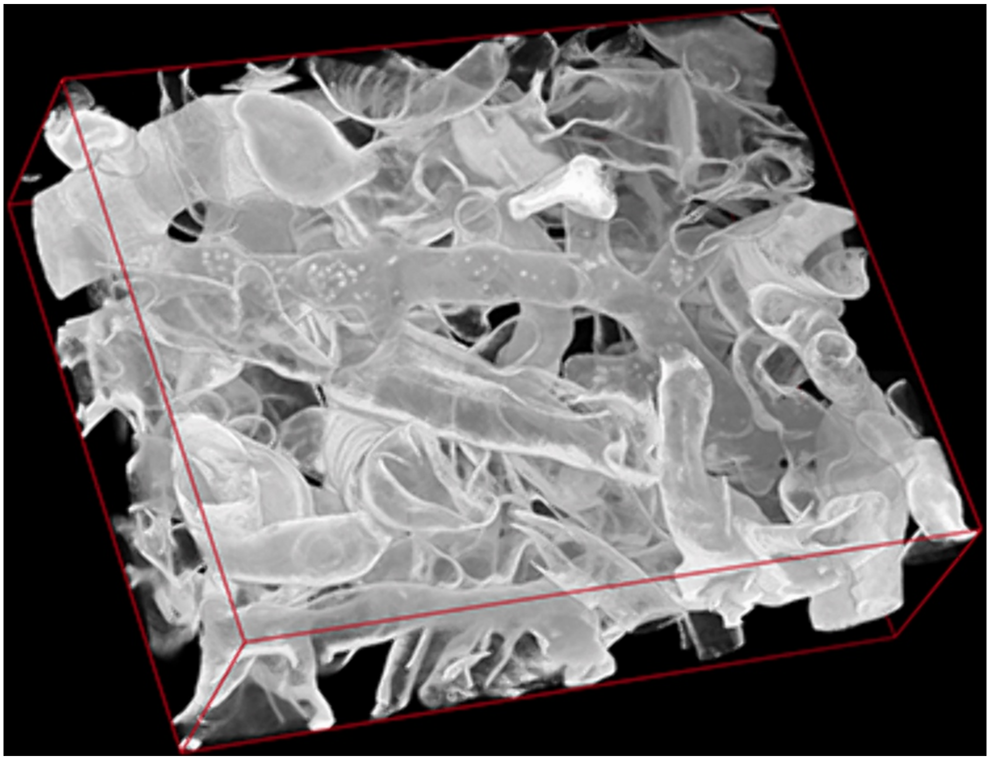

**Figure S9: 3D visualization of FIBSEM data**. A 3D view of the 375 images acquired from the wild-type dataset and aligned, prior to volume segmentation

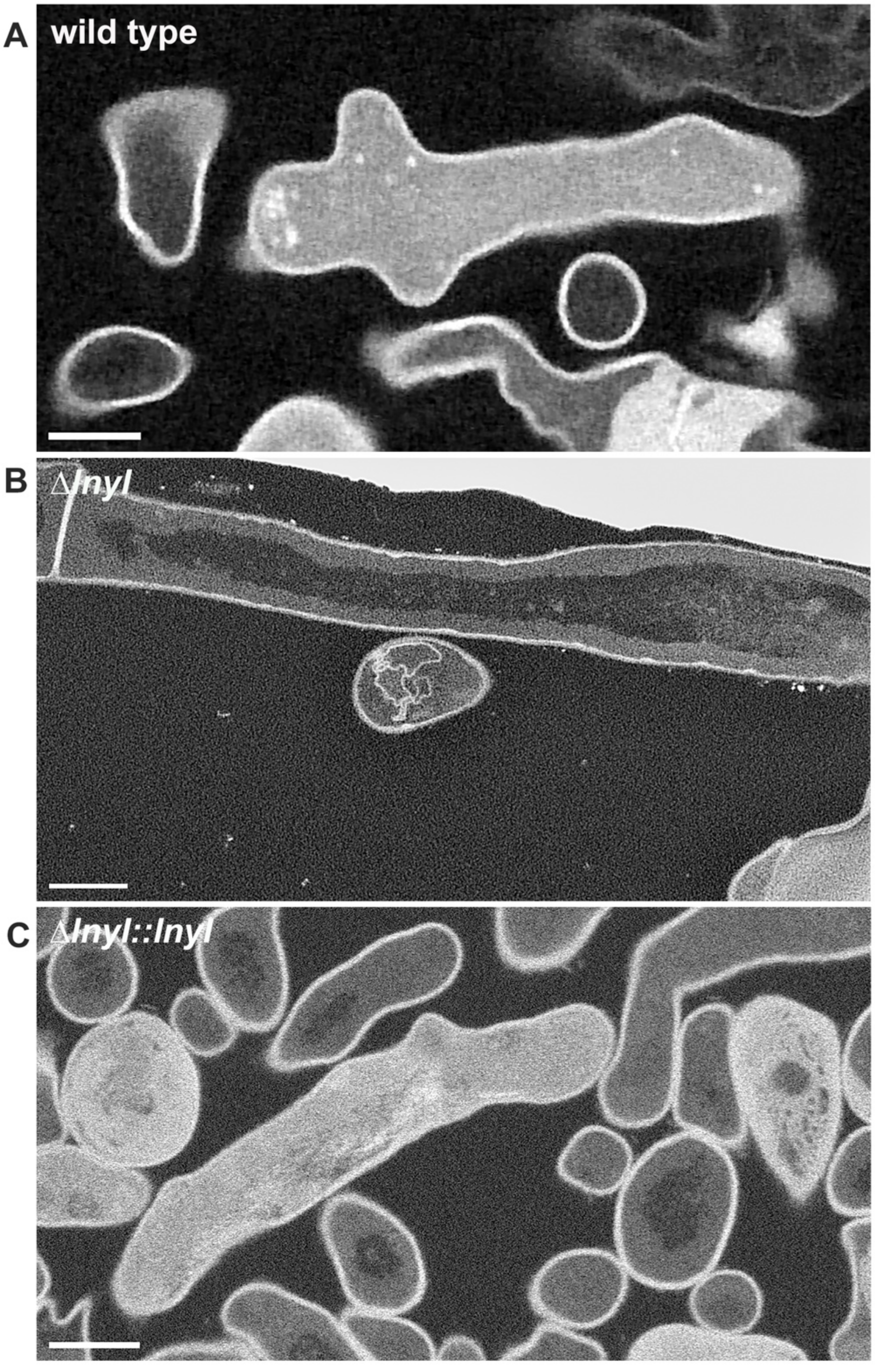


**Figure S10: Two-dimensional view of individual FIB-SEM images from serial imaging datasets**. 2D view of the selected filaments from (A) wild-type, (B) Δ*lnyI*, and (C) Δ*lnyI::lnyI S*. Mg1 strains. The scale bar = 1 μm.

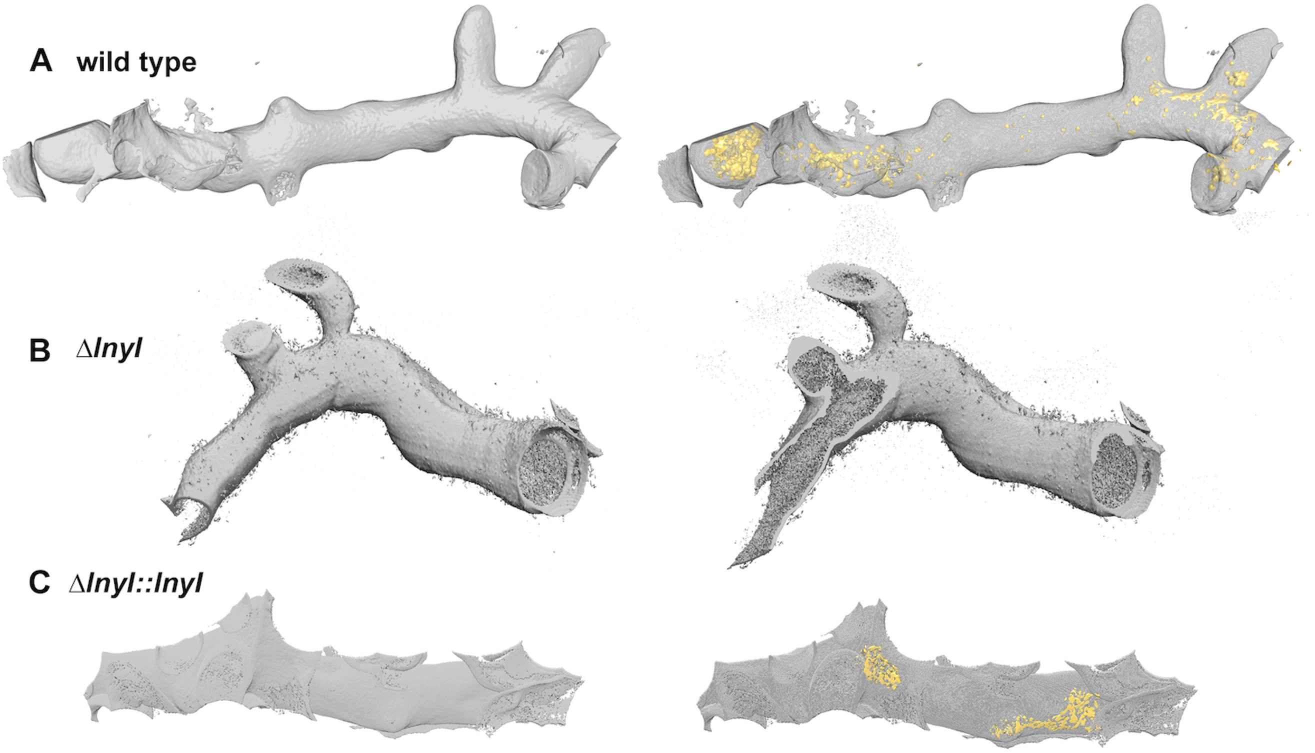


**Figure S11: Independent volume representation of hyphae and their subcelluar densities**. Additional examples of three-dimensional reconstructions for (A) wild type, (B) ∆*lnyI*, and (C) ∆*lnyI::lnyI S*. Mg1 strains. Images on the left are original opacity following segmentation and image processing. Images on the right are the same hyphae where the cellular surface is rendered partially transparent to reveal the location and forms of the subcellular compartments. The compartments are absent in the ∆*lnyI* strain.

**Table S1. Linearmycin signal overlap with membrane-intense regions**

| Panel No. | FM4-64 signal * | Linearmycin signal overlap | % signal overlap | Nile red signal | Linearmycin signal overlap | % signal overlap |
|---|---|---|---|---|---|---|
| 1 | 19 | 15 | 79 | 25 | 21 | 84 |
| 2 | 17 | 14 | 82 | 28 | 27 | 96 |
| 3 | 28 | 23 | 82 | 50 | 45 | 90 |
| 4 | 25 | 20 | 80 | 50 | 46 | 92 |
| 5 | 14 | 10 | 71 | 34 | 30 | 88 |

* Total number of stained membranous regions counted in the hyphae for each panel.

**Table S2: Bacterial strains used in the study**

| Strain | Genotype | Source |
|---|---|---|
| PSK0558 | *Streptomyces* sp Mg1 | (9) |
| PSK0207 | *Bacillus subtilis* NCIB 3610 | (9) |
| PDS 0755 | *Streptomyces sp*. Mg1 Δ*lnyI::apr* | (9) |
| PDS0794 | *Streptomyces sp*. Mg1 Δ*lnyI::lnyI* (apr, hyg) | (9) |
| PDS1332 | *Streptomyces sp*. Mg1 Δ*lnyI::lnyI-ypet* (apr, hyg) | This study |
| PDS1340 | *Streptomyces sp* Mg1 *attB*:: PrpsL-Ypet | This study |

**Table S3. Primer sequences used in this study**

| Primer name | Primer sequence (5'-3') |
|---|---|
| pOOB40-FP | atcgaattcgtaatcatgtcatagctgtttcctgtg |
| pOOB40-RP | atcgcgcgcggccgcggatc |
| lnyI-FP | gatccgcggccgcgcgcgat |
| lnyI-RP | gctcccgccgcctcccggcctcgggacggc |
| lnyI- yfp-FP | ggaggcggcgggagcatggtctccaagggc |
| pOOB40-yfp-RP | gacatgattacgaattcgattcacttgtacagctcgttc |
| BCH0067 | gatgcggtattttctccttacgc |
| BCH0068 | ggttttcccagtcacgacgtt |
| BCH0070 | aaggagaaaataccgcatctcacttgtacagctcgttcatgccc |
| BCH0071 | gtcgtgactgggaaaaccaggaggtccatatggtctccaagggc |
| BCH0072 | catatggacctccttacgtctccgtcgtctactcgagc |
| BCH0073 | ggattcggatccgccctgcaggcggaagtcag |

**Table S4: Parameters and conditions for FIB sectioning and SEM image acquisition for *S*. Mg1 strains**.

| Parameters | Wild type | Δ*lnyI* | Δ*lnyI* ::*lnyI* |
|---|---|---|---|
| FIB voltage | 30 kV | 30 kV | 30 kV |
| FIB current | 500 pA | 400 pA | 400 pA |
| SEM voltage | 2 keV | 2 keV | 2 keV |
| SEM current | 100 pA | 260 pA | 260 pA |
| Slice thickness | 10 nm | 10 nm | 10 nm |
| SEM Detector | T1 (A+B) | ICD | ICD |
| SEM Horizontal field width (HFW) | 15.36 μm | 15.2 μm | 15.37 μm |
| Dwell time | 4 μs | 4 μs | 4 μs |
| SEM pixel size | 5 x 5 $nm^2$ | 5 x 5 $nm^2$ | 5 x 5 $nm^2$ |
| Total no. of images | 325 | 301 | 578 |

**See links for Movies 1 – 3**

**M1**

**M2**

**M3**
